# Congenital glaucoma with anterior segment dysgenesis in individuals with biallelic *CPAMD8* variants

**DOI:** 10.1101/297077

**Authors:** Owen M Siggs, Emmanuelle Souzeau, Deepa A Taranath, Tiger Zhou, Andrew Dubowsky, Shari Javadiyan, Angela Chappell, Andrew Narita, James E Elder, John Pater, Jonathan B Ruddle, James EH Smith, Lisa S Kearns, Sandra E Staffieri, Alex W Hewitt, David A Mackey, Kathryn P Burdon, Jamie E Craig

## Abstract

**Purpose:** Congenital glaucoma is a significant cause of irreversible blindness. In some instances glaucoma is associated with developmental abnormalities of the ocular anterior segment, which can impair drainage of aqueous humor, leading to an increase in intraocular pressure.

**Methods:** Genome sequencing was performed on a parent-proband congenital glaucoma trio, with exome sequencing of 79 additional individuals with suspected primary congenital glaucoma.

**Results:** We describe a unique ocular anterior segment dysgenesis associated with congenital glaucoma in four individuals from three unrelated families. In each case, disease was associated with compound heterozygous variants in *CPAMD8*, a gene of unknown function recently associated with ocular anterior segment dysgenesis, myopia, and ectopia lentis. *CPAMD8* expression was highest in neural crest-derived tissues of the adult anterior segment, suggesting that *CPAMD8* variation may cause malformation of key drainage structures and the development of high intraocular pressure and glaucoma.

**Conclusions:** This study reveals a unique genetic cause of childhood glaucoma, and expands the phenotypic spectrum of *CPAMD8*-associated ocular disease.

## Introduction

Congenital glaucoma is a significant cause of childhood blindness and results from developmental defects of the trabecular meshwork and Schlemm’s canal, both of which are involved in the regulation of aqueous humor outflow controlling intraocular pressure (IOP). Increased IOP is a major risk factor for optic disc cupping and retinal nerve fibre layer degeneration: the major hallmarks of glaucoma. Other common features of congenital glaucoma include buphthalmos, corneal clouding, corneal edema, and breaks in Descemet’s membrane (Haab striae). Congenital glaucoma can be primary (PCG) in which there is no other apparent ocular abnormality, or secondary, in which other ocular features are present. Secondary congenital glaucoma is often caused by anterior segment dysgenesis (ASD), a heterogeneous group of ocular developmental disorders affecting the iris, cornea, lens, and/or aqueous humor outflow structures. Associated defects include hypoplasia of the iris stroma, corectopia, pseudopolycoria, posterior embryotoxon, abnormal iridocorneal angle, iridocorneal adhesions, corneal opacity, and corneal vascularisation. The most common ASD presentation is Axenfeld-Rieger anomaly, known as Axenfeld-Rieger syndrome (ARS) when associated with systemic features. Individuals with ARS have a 50-60% lifetime risk of developing glaucoma, which can occur as early as the first year of life (Shields 1983; Souzeau 2017).

Three genes have been associated with PCG: *CYP1B1* ^1^, *LTBP2* ^2,3^ and *TEK* ^4^. Depending upon the population, biallelic variants in *CYP1B1* account for as few as 15% to as many as 100% of PCG cases ^5,6^, with a prevalence of 22% among Australian cases ^7^. *LTBP2* variants are a rare cause of autosomal recessive congenital glaucoma and are often associated with megalocornea, spherophakia, and ectopia lentis ^8^. Heterozygous loss-of-function *TEK* variants are also a rare cause of PCG with incomplete penetrance ^4^. While heterozygous variants in *FOXC1* ^9^ or *PITX2* ^10^ account for 40-60% of ARS, *FOXC1* variants may also be associated with childhood glaucoma with mild features of ASD (Siggs et al., unpublished). Similarly, *PAX6* variants cause aniridia ^11^, and in rare instances can be associated with glaucoma ^12^.

Recently, a unique form of ASD was associated with biallelic variants in *CPAMD8* ^13^ (Anterior segment dysgenesis 8, OMIM 617319). Unlike other ASD cases, none of the four original cases with biallelic *CPAMD8* variants had glaucoma; only a single case was reported to have developed ocular hypertension at 49 years of age, suspected to have been exacerbated by ectopia lentis and retinal surgery ^13^. A homozygous variant in *CPAMD8* has also been associated with Morgagnian cataract in Red Holstein Friesian cattle, with glaucoma observed in one of eight eyes examined, and associated with uveitis and retinitis in the same eye ^14^.

Here we investigated genetic variation in a cohort of patients with a suspected diagnosis of PCG. Compound heterozygous *CPAMD8* variants were identified in four affected individuals from three families, each with a unique combination of anterior segment abnormalities and congenital glaucoma.

## Material and Methods

### Human subjects

Patients and family members were recruited via the Australian & New Zealand Registry of Advanced Glaucoma ^15^ and provided written informed consent under protocols approved by the Southern Adelaide Clinical Human Research Ethics Committee. The study was conducted in accordance with the revised Declaration of Helsinki and followed the National Health and Medical Research Council statement of ethical conduct in research involving humans. Clinical details were obtained from the treating ophthalmologist.

### DNA sequencing and analysis

DNA was prepared from whole blood. Genomes were sequenced on the Illumina Hiseq X platform as previously described ^16^. Exome capture was performed using the Agilent SureSelect system (v4 or v5) and sequenced on an Illumina HiSeq (2000 or 2500). Raw reads were aligned to the hg19 reference using BWA-MEM (v0.7.12), and sorted and duplicate-marked with Picard (v1.13). Local indel realignment and base quality score recalibration was performed using GATK (v3.4.0). Variants were recalibrated using the GATK Variant Quality Score Recalibrator (VQSR). Finally, VCF files were annotated with ANNOVAR using RefSeq, gnomAD (r2.0.2), 1000 Genomes (Phase 3), NHLBI-ESP project (v2), ClinVar (20170905), and dbNSFP (v3.3a) databases. The Combined Annotation Dependent Depletion (CADD), Sorting Intolerant From Tolerant (SIFT), and PolyPhen-2 HumVar tools were used to predict the effect of sequence variants on protein function. Protein sequences were aligned with Clustal Omega and BOXSHADE.

Confirmation of variants identified through genome and exome sequencing and segregation studies were performed by a National Association of Testing Authorities (NATA)-accredited laboratory (SA Pathology, Flinders Medical Centre, Adelaide, Australia) using bi-directional capillary sequencing of the relevant PCR-amplified *CPAMD8* region. Primer sequences are available upon request. PCR products were sequenced and base called on the Applied Biosystems 3130*xl* Genetic Analyzer (ThermoFisher Scientific). Detection of sequence variants was performed with the aid of Mutation Surveyor v4.0 (SoftGenetics LLC), with trace files assembled against the *CPAMD8* (NM_015692.2) hg19 reference.

### Tissue dissection and RNA preparation

Cadaveric human eyes with no known ocular disease and otherwise unsuitable for corneal transplantation were obtained from the Eye Bank of South Australia. Tissue dissection was performed under light microscope with a mean post-mortem time of 9.7 ± 5.3 hours. Tissues from the corneal epithelium, corneal stroma, corneal endothelium, trabecular meshwork (TM), pars plicata of the ciliary body, retina, optic nerve head, and optic nerve were collected and fixed in RNAlater solution for approximately 5 days prior to storage at -80°C. A standard Trizol extraction protocol was used for RNA isolation. RNA extracted from the pars plicata was passed through a Genomic-tip 20/G (Qiagen) as per the manufacturer’s instructions to remove excess melanin. RNA quality was assessed using the Agilent Bioanalyzer 2100 RNA 6000 Nano Assay (mean RIN = 6.5 ± 1.8). The Qubit® 2.0 Fluorometer (Thermo Fisher Scientific) was used to quantify RNA.

### RNA sequencing and analysis

RNA libraries were prepared using the Bioo Scientific^®^ NEXTflex™ Rapid Directional mRNA-Seq Kit Bundle with RNA-Seq Barcodes and poly(A) beads. Total RNA from each tissue sample (250 ng) was prepared for individual sequencing. Sequencing was performed on an Illumina NextSeq^®^ 500 using High Output v2 Kit (75 cycles) following sample enrichment with 16 cycles of PCR. Phi X DNA was spiked into each run pool. Trimgalore v0.4.0 was used to trim sequence adaptors and low quality bases (Phred scores < 28). Remaining reads were aligned to the human genome (GRCh38 assembly) using TopHat v2.1.1, allowing for 2 mismatches per read and no ambiguous reads. Uniquely aligned reads were annotated using the union model in HTSeq-count v0.6.0 with Ensembl version 84 human gene ID. Trimmed mean of M-values (TMM) was used to normalize count data, followed by calculation of differential expression in EdgeR v3.10.2. Benjamini Hochberg adjustment was applied for false discovery rate correction.

## Results

To investigate the underlying genetic cause of congenital glaucoma, we sequenced the genomes of an affected proband and his unaffected parents (Family 1, Figure 1A). Based on a suspected autosomal recessive mode of inheritance, we initially focussed on all genes with rare homozygous or compound heterozygous variants (gnomAD allele frequency < 0.001) present in the proband (which included *SCAF1, TTN, TXND8*, and *CPAMD8*). Only one of these genes (*CPAMD8*) had previously been associated with ocular disease, and furthermore was known to cause an autosomal recessive ASD ^13^. The two heterozygous *CPAMD8* variants present in the proband (p.(Arg1135Ter), p.(Pro1188Leu)) were confirmed to be *in trans*, with one variant present in each parent (Figure 1A).

**Figure 1.**
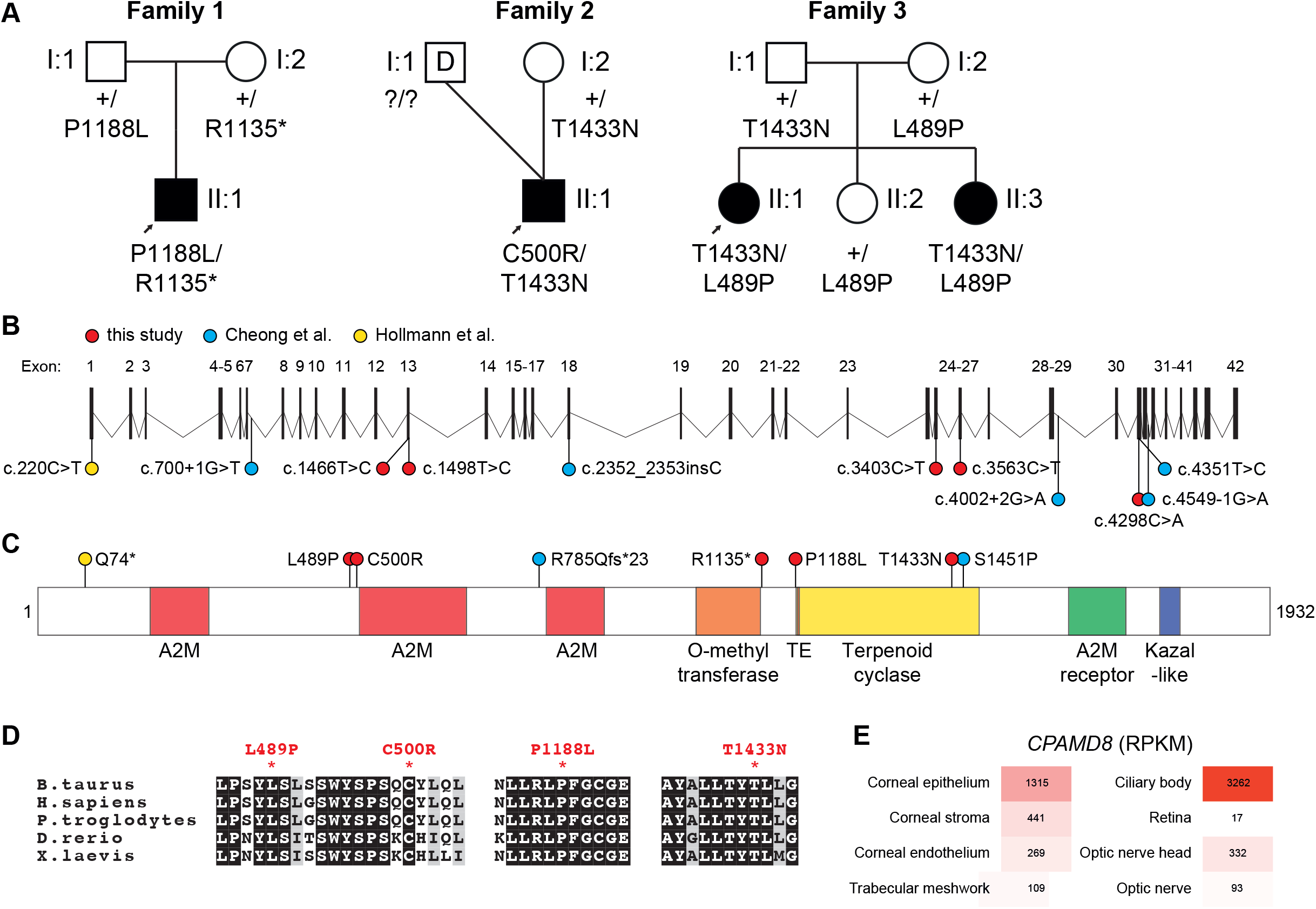
Congenital glaucoma with anterior segment dysgenesis in individuals with biallelic *CPAMD8* variants. A. Pedigrees of the three families described in this study. In Family 2, the proband was conceived via an anonymous sperm donor. Round symbols indicate females; square symbols, males; black symbols, congenital glaucoma; arrow, proband; D, sperm donor; +, reference allele. B. *CPAMD8* locus and location of disease-associated variants described in this study, and others ^13,14^. C. CPAMD8 protein schematic and domain structure, showing location of disease-associated missense, nonsense, and frameshift variants (splice variants not shown). A2M, α2-macroglobulin domain; TE, thioester site. D. CPAMD8 protein sequence alignments across multiple vertebrate species. Residues of interest are highlighted for variants p.(Leu489Pro), p.(Cys500Arg), p.(Pro1188Leu) and p.(Thr1433Asn). E. *CPAMD8* expression in adult donor ocular tissue. RPKM, reads per kilobase per million mapped reads.

Given the phenotypic overlap between ASD and congenital glaucoma (Siggs et al., unpublished), we expanded our search for *CPAMD8* variants across a further 79 exomes from individuals with suspected PCG. We identified rare (gnomAD allele frequency < 0.001) compound heterozygous variants in an additional three patients with congenital glaucoma from two unrelated families (Figure 1A-C), consistent with suspected autosomal recessive inheritance in both families. In Family 2, the proband (II:1) was conceived via anonymous sperm donation, with the unaffected mother confirmed to carry one heterozygous variant. In Family 3, two affected sisters (II:1 and II:3), but not a third unaffected sister (II:2), carried the same pair of compound heterozygous variants, with one variant carried by each unaffected parent (Figure 1A).

All five *CPAMD8* variants identified were either rare or absent in gnomAD, at a frequency consistent with a rare pediatric recessive disease (Table 1), and no homozygotes were observed. All identified *CPAMD8* variants were predicted to be damaging by the CADD, PolyPhen-2 and/or SIFT algorithms (Table 1). Alignment of vertebrate CPAMD8 protein sequences demonstrated strong conservation at the position of each of the missense variant residues (Figure 1D). Families 2 and 3 were not known to be related but both are of Scottish ancestry, and shared a single variant (p.(Thr1433Asn)) found only in the non-Finnish European cohort of gnomAD (5/125954 alleles).

**Table 1.** *CPAMD8* variants associated with anterior segment dysgenesis and congenital glaucoma

| Family | gDNA (hg19) | cDNA<br>(NM_015692) | exon | protein (NP_056507) | gnomAD AF | CADD_<br>phred | PP2_HVAR | SIFT |
| --- | --- | --- | --- | --- | --- | --- | --- | --- |
| **1** | g.19:17038927G>A | c.3403C>T | 25 | p.(Arg1135Ter) | 10/276872<br>(0.00003612) | 39 | . | . |
|  | g.19:17036131G>A | c.3563C>T | 26 | p.(Pro1188Leu) | 4/246198<br>(0.00001625) | 33 | 1; D | 0.031;<br>D |
| **2** | g.19:17015130G>T | c.4298C>A | 32 | p.(Thr1433Asn) | 5/276098<br>(0.00001811) | 25.6 | 0.955; D | 0.008;<br>D |
|  | g.19:17100491A>G | c.1498T>C | 13 | p.(Cys500Arg) | . | 26.2 | 0.997; D | 0.006;<br>D |
| **3** | g.19:17015130G>T | c.4298C>A | 32 | p.(Thr1433Asn) | 5/276098<br>(0.00001811) | 25.6 | 0.955; D | 0.008;<br>D |
|  | g.19:17100523A>G | c.1466T>C | 13 | p.(Leu489Pro) | 1/245962<br>(0.000004066) | 26.4 | 0.963; D | 0.211;<br>T |

Position of each predicted pathogenic variant with the human genome, consensus transcript, and protein described using HGVS nomenclature (version 15.11). Frequencies of each allele in the gnomAD database is shown, as are CADD, PolyPhen-2, and SIFT scores with predictions of deleteriousness. D, deleterious; T, tolerated.

*CPAMD8* is known to be expressed in the anterior segment of the developing human eye, with high expression of transcript at 22 weeks’ gestation in the lens, iris, and cornea ^13^. We further assessed *CPAMD8* expression in ocular tissue from recently deceased adult donors, identifying a peak of expression in the ciliary body and corneal epithelium (Figure 1E).

The phenotype of each affected family member is detailed in Figure 2 and Table 2. The proband of Family 1 (II:1) was diagnosed with bilateral buphthalmos and congenital glaucoma at birth. He required multiple operations to control his elevated IOP and glaucoma (bilateral trabeculotomies and trabeculectomies, and a glaucoma drainage tube implant in the right eye), along with topical eye drops. Other ocular features included high myopia (due to increased axial length secondary to buphthalmos), iris stromal hypoplasia, microphakia, scleromalacia, prominent iris vessels, and pendular nystagmus. He also had ectopia lentis, corectopia, and iris transillumination, all of which were suspected to be consequences of surgery. Both parents had a normal eye examination with no features of ASD.

**Figure 2.**
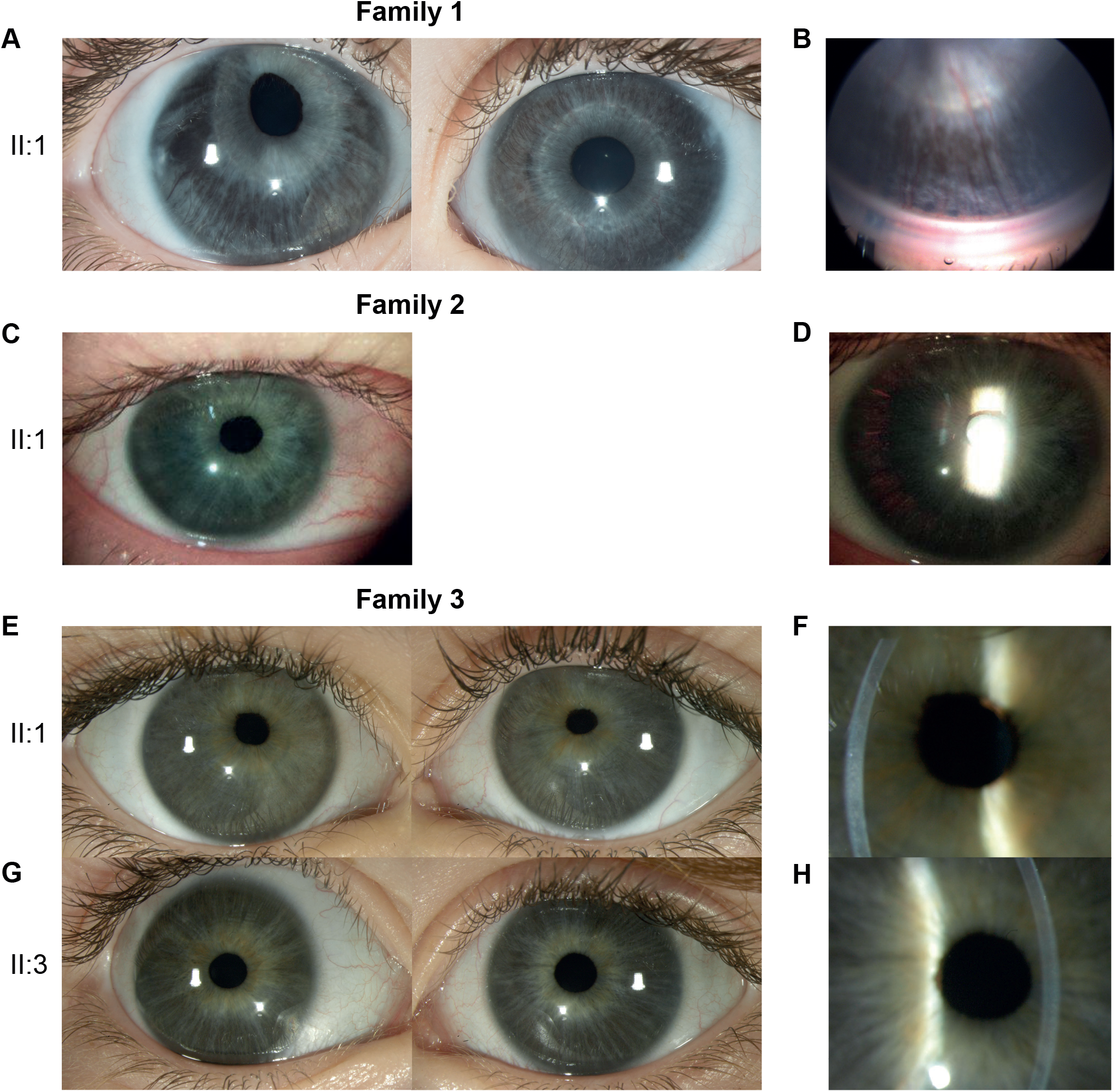
Anterior segment dysgenesis phenotypes in individuals with biallelic *CPAMD8* variants. A, C, E, G. Clinical photographs of the right eye (RE, left panels) and the left eye (LE, right panels) of the specified individuals. Family 1 (A,B): superonasal corectopia, suspected to be secondary to or exacerbated by trabeculotomy surgery (A, RE), iris stromal hypoplasia (A, both eyes), and prominent iris vessels with open angles on Retcam gonioscopy (B, LE). Family 2 (C,D): mild superonasal corectopia (C, RE), mild iris stromal hypoplasia (C, RE), and iris transillumination (D, RE). Family 3 (E-H): superonasal corectopia (E, both eyes; G, LE), iris stromal hypoplasia (G, both eyes), and mild ectropion uveae (F, RE; H, LE).

**Table 2.**
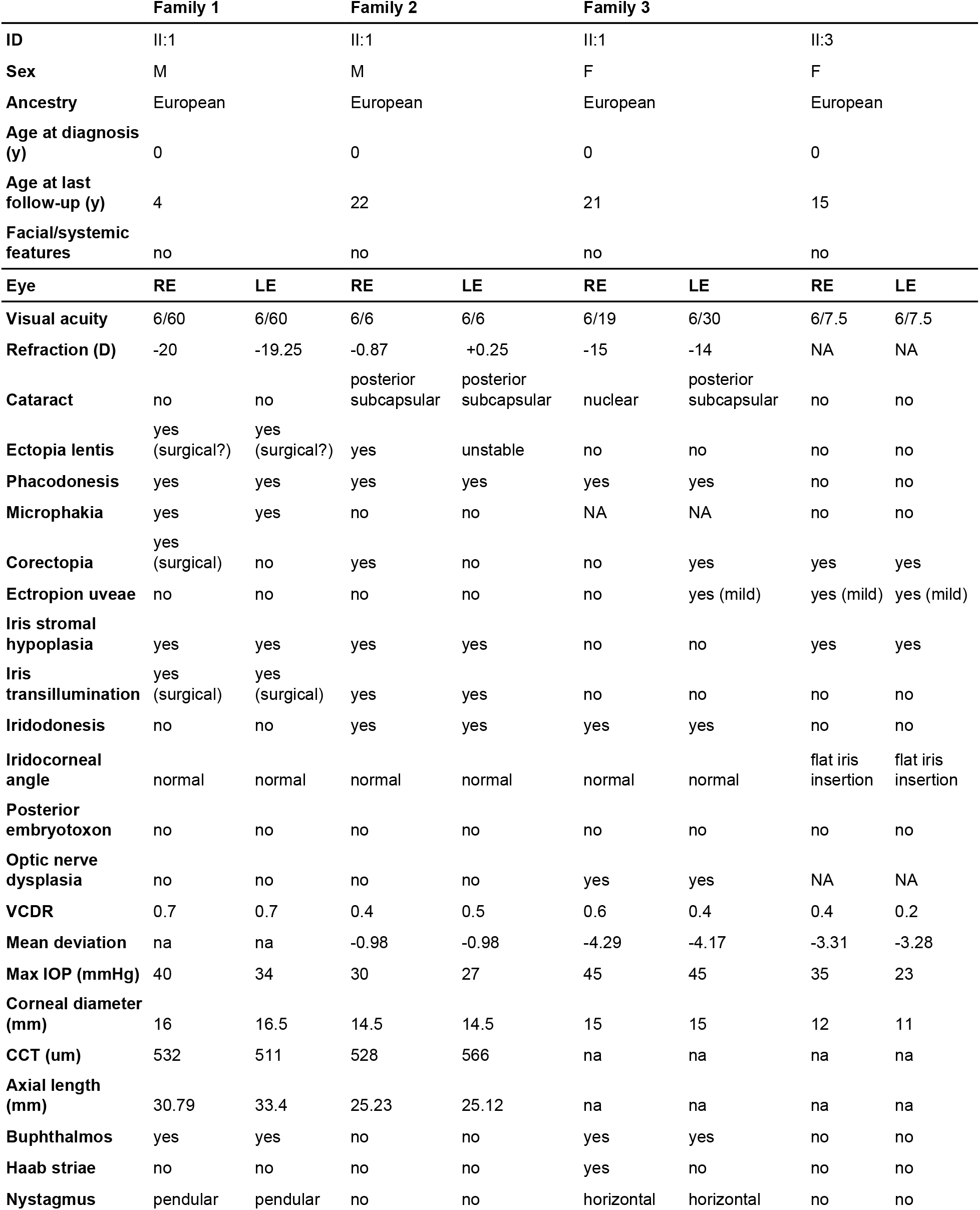
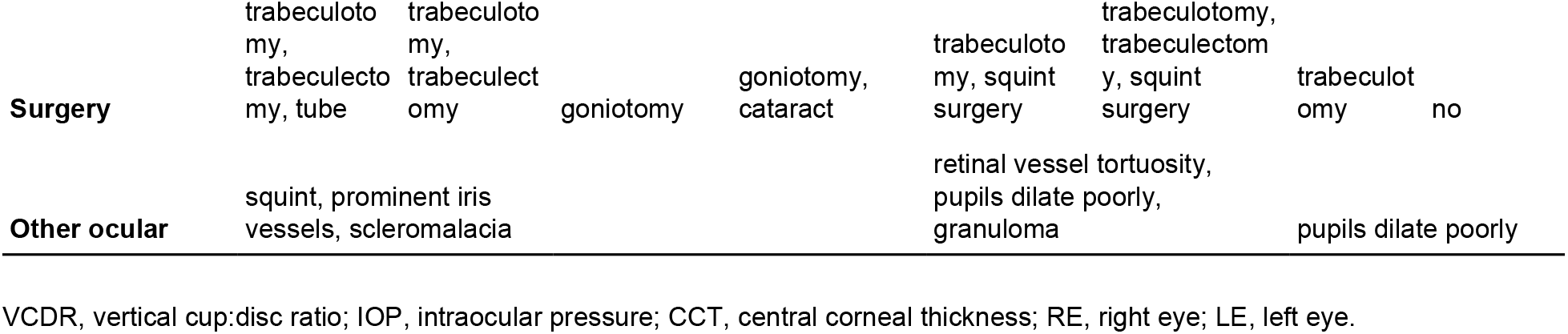
Clinical details of four individuals with compound heterozygous *CPAMD8* variants

The proband of Family 2 (II:1) was diagnosed with congenital glaucoma at birth, with bilateral megalocornea, nystagmus, and elevated IOP. He had bilateral goniotomies at 3 days of age to control IOP, which remained controlled ever since. He developed bilateral posterior subcapsular cataracts at 22 years of age, and had a left cataract removed (due to megalocornea he remained aphakic in that eye). Other ocular features included iris stromal hypoplasia, corectopia, iridodonesis, phacodonesis, and ectopia lentis. His mother had a normal eye examination. He was conceived through an anonymous sperm donor; therefore, no clinical details were available for his biological father. He had an unaffected younger brother who was conceived through a different sperm donor.

The proband of Family 3 (II:1) was diagnosed with bilateral congenital glaucoma at birth. She had trabeculotomy in the right eye at two weeks of age, followed by trabeculectomies in both eyes at three weeks of age. Her IOPs had been well maintained ever since and she was not on any treatment at her last follow-up (at 21 years of age). She had high myopia, iris stromal hypoplasia, corectopia, phacodonesis, iridodonesis, ectropion uveae, and nystagmus. She developed nuclear and posterior subcapsular cataracts at 6 years of age, which had not required surgery at last follow-up. Her sister (II:3) was diagnosed with unilateral PCG at birth, and underwent a right trabeculotomy at one week of age, with well controlled IOP since this procedure. Other ocular features included iris stromal hypoplasia, corectopia, and corectopia. Both parents and a third sibling had normal eye examinations.

Clinical data summarised across all four affected individuals revealed striking similarities to previously described cases of *CPAMD8*-associated ASD (Figure 2, Table 2) ^13^. All cases were noted to have abnormalities of the iris (4/4), including corectopia (4/4), stromal hypoplasia (3/4), and mild ectropion uveae (2/4) (Figure 2). Cataracts were present in both individuals older than 20 years, with cataract surgery complicated by megalocornea in the proband of Family 2. All four cases described here required surgery to control glaucoma and IOP: three had well controlled IOP and glaucoma after surgery in early infancy, while the proband of Family 1 required multiple surgical procedures (and was still on topical treatment at 4 years of age) (Table 2).

## Discussion

We provide evidence that biallelic *CPAMD8* variation represents a new genetic cause of congenital glaucoma. Along with its unique ocular anterior segment findings, this extends the spectrum of phenotypes associated with *CPAMD8* variants. Within an Australian cohort of suspected PCG probands, biallelic *CPAMD8* variants accounted for at least 3.8% of individuals (3/80), or 3.0% (3/100) when including individuals explained by variants in *CYP1B1*. This rate is similar to the frequency associated with heterozygous loss-of-function variants in *TEK* ^4^, and *FOXC1* (Siggs et al., unpublished). Other ASDs, notably ARS caused by *FOXC1* and *PITX2* variants, have a well-defined association with glaucoma ^17^.

Ascertainment bias makes it difficult to accurately estimate glaucoma penetrance in *CPAMD8*-associated ASD. Within the limits of available patient data (four known *CPAMD8* cases with glaucoma, plus four without glaucoma), we estimate glaucoma penetrance in *CPAMD8*-related ASD to be ~50% (4/8), which is similar to more reliable estimates in ARS ^17,18^. More commonly associated ocular features include corectopia (8/8), iris stromal hypoplasia (6/8), iris transillumination defects (5/8), iridodonesis (5/8), and ectopia lentis (5/8). Other features commonly associated with ASD (corneal opacity and posterior embryotoxon) were absent. Cataracts were present in 6/6 individuals above 20 years of age.

The ocular phenotype of ARS is clearly distinct from *CPAMD8*-associated ASD, suggesting that the two entities may cause glaucoma via different mechanisms. The conspicuous absence of a *CPAMD8* ortholog in rodent genomes makes this gene difficult to study in model organisms, although some inference can be made from expression studies. *CPAMD8* is consistently expressed in tissues of neural crest origin ^19^ including the lens, iris and cornea *in utero* ^13^, and the ciliary body and corneal epithelium in adult eyes. This pattern is consistent with hypoplasia of the iris stroma in patients with *CPAMD8* variants, while neuroectoderm-derived pigmented epithelial cells develop normally. Other key drainage structures in the eye, including the trabecular meshwork and Schlemm’s canal, are also derived from neural crest progenitors, and may be hypoplastic or otherwise dysfunctional in the absence of CPAMD8. But unlike other ASDs defined by neural crest dysfunction ^20^, *CPAMD8* deficiency does not appear to cause other neural crest-associated malformations involving dental, craniofacial, nor aortic structures.

Initially suspected to play a role in immunity, the physiological function of *CPAMD8* has been elusive, particularly given its absence from rodent genomes ^21^. CPAMD8 (complement component 3- and pregnancy zone protein-like alpha-2-macroglobulin domain-containing protein 8) is one of eight members of the complement 3/α2-macroglobulin (C3/α2M) family reported in humans ^21^. Other C3/α2M family members include the complement components C3, C4A, C4B, and C5; α2M; pregnancy zone protein; and CD109. C3, C4A, C4B, C5, and α2M proteins are all abundant in plasma, while CD109 is GPI-anchored cell-surface molecule, and CPAMD8 is membrane-associated.

In conclusion, our findings expand the phenotypic spectrum associated with biallelic *CPAMD8* variants, and extend the overlap between congenital glaucoma and ASD (Siggs et al., unpublished)^17^: highlighting the challenges associated with clinically distinguishing the two. Molecular diagnostics will play an increasingly important role in the diagnosis and management of childhood and congenital glaucoma.

## Acknowledgements

We thank Carly Emerson for photography. This project was supported by the Channel 7 Children’s Research Foundation, and the Australian National Health & Medical Research Council (NHMRC, Centres of Research Excellence Grant 1023911). KPB and AWH were supported by an NHMRC Senior Research Fellowship, and JEC by an NHMRC Practitioner Fellowship. The Centre for Eye Research Australia (CERA) received Operational Infrastructure Support from the Victorian Government.

